# Synergistic Hypoxia and Apoptosis Conditioning Unleashes Superior Mesenchymal Stem Cells Efficacy in Acute Graft-versus-Host-Disease

**DOI:** 10.1101/2024.04.11.588248

**Authors:** Mohini Mendiratta, Meenakshi Mendiratta, Shuvadeep Ganguly, Sandeep Rai, Ritu Gupta, Lalit Kumar, Sameer Bakhshi, Vatsla Dadhwal, Deepam Pushpam, Prabhat Singh Malik, Raja Pramanik, Mukul Aggarwal, Aditya Kumar Gupta, Rishi Dhawan, Tulika Seth, Manoranjan Mahapatra, Baibaswata Nayak, Thoudam Debraj Singh, Sachin Kumar Singla, Mayank Singh, Chandra Prakash Prasad, Hridayesh Prakash, Sujata Mohanty, Ranjit Kumar Sahoo

**Affiliations:** Department of Medical Oncology, Dr. B. R. Ambedkar Institute Rotary Cancer Hospital, All India Institute of Medical Sciences, New Delhi, India-110029; Stem Cell Facility (DBT-Centre of Excellence for Stem Cell Research), All India Institute of Medical Sciences, New Delhi, India-110029; Laboratory Oncology Unit, Dr. B. R. Ambedkar Institute Rotary Cancer Hospital, All India Institute of Medical Sciences, New Delhi, India-110029; Department of Obstetrics and Gynecology, All India Institute of Medical Sciences, New Delhi, India-110029; Department of Hematology, All India Institute of Medical Sciences, New Delhi, India-110029; Department of Pediatrics Oncology, All India Institute of Medical Sciences, New Delhi, India-110029; Department of Gastroenterology and Human Nutrition, All India Institute of Medical Sciences, New Delhi, India-110029; Amity Centre for Translational Research, Amity University, Sector 125, Noida, India 201313

**Keywords:** Mesenchymal Stem Cells, Apoptosis, Hypoxia, Immunomodulation, Acute Graft-versus-Host-Disease, Wharton’s Jelly, Bone marrow

## Abstract

Mesenchymal stem cells (MSCs) have emerged as promising candidates for immune modulation in various diseases that are associated with dysregulated immune responses like Graft-versus-Host-Disease (GVHD). MSCs are pleiotropic and the fate of MSCs following administration is a major determinant of their therapeutic efficacy. In this context, we here demonstrate that hypoxia preconditioned apoptotic MSCs [bone marrow (BM), Wharton’s Jelly (WJ)] bear more immune programming ability in a cellular model of acute Graft-versus-Host-Disease (aGVHD). To this purpose, we programmed MSCs by exposing them to hypoxia and inducing apoptosis both sequentially as well as simultaneously. Our findings demonstrated that WJ MSCs that were conditioned with indicated approaches simultaneously induced the differentiation of CD4^+^T-cell towards Tregs, enhanced Th2 effector, and concomitantly mitigated Th1 and Th17, with polarization of M1 effector macrophages towards their M2 phenotype, and more interestingly enhanced efferocytosis by macrophages indicated Th2 programming ability of MSCs programmed by conjunctional approaches Overall, our study highlights the potential of WJ-MSCs conditioned with hypoxia and apoptosis concurrently, as a promising therapeutic strategy for aGVHD and underscores the importance of considering MSC apoptosis in optimizing MSCs-based cellular therapy protocols for enhanced therapeutic efficacy in aGvHD.

**GRAPHICAL ABSTRACT:** 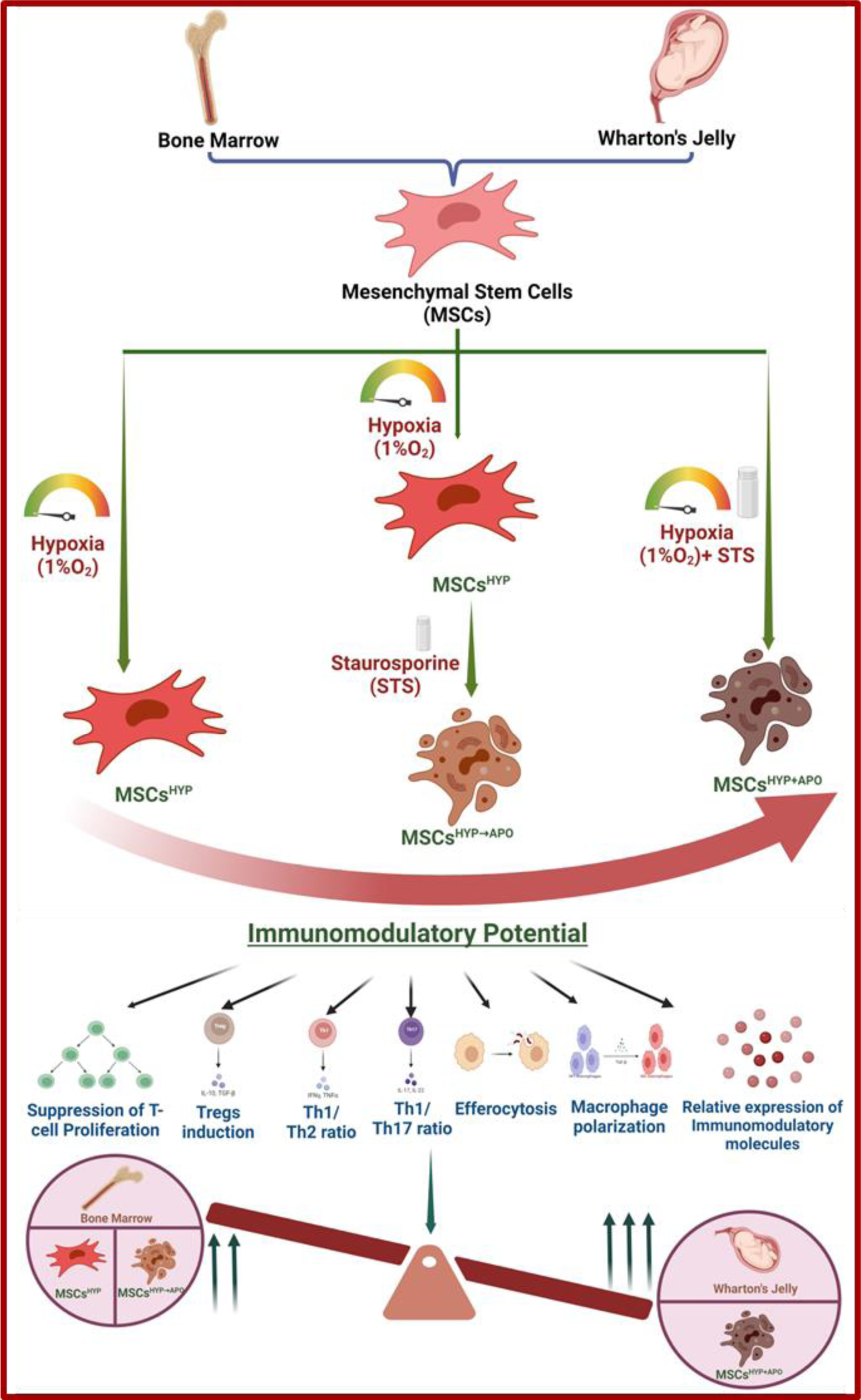

## INTRODUCTION

Mesenchymal stem cells (MSCs) exert their immunoregulatory effects in Graft-versus-Host-Disease (GVHD) (1,2) through the secretion of various soluble factors such as chemokines, cytokines, and extracellular vesicles/exosomes (3), resulting in modulation of the immune response by suppressing the activation of immune cells (T-cell, natural killer cells, dendritic cells, B-cell) in an inflammatory milieu (4,5).

Several preclinical studies reported that transplanted MSCs have a limited lifespan in recipients as they undergo apoptosis by various immune effector cells (cytotoxic T-cell, natural killer cell, granulocytes) (6–8) within the host circulation and apoptotic MSCs are subsequently efferocytosed by macrophages which in turn licensed to disseminate soluble factors to regulate the activated immune response (9). Therefore, this mechanism has been explored in a few studies by either inducing or inhibiting the apoptosis in MSCs by silencing the apoptotic effector molecules (BAK and BAX) in the MSCs and validating their therapeutic potential (10) and demonstrating that there was a reduction in their immunomodulatory capabilities in a viable form when administered in an asthmatic mice model, thereby potentiating the need for MSCs for undergoing apoptosis to exert their functional effect (10–12). Moreover, these apoptotic MSCs have been found to possess the ability to induce immunosuppressive effects in animal models of GVHD and various organ injuries like lung, liver, and spleen (6,13,14), suggesting that phagocytes are mediators of MSC-induced adaptive responses (15).

Moreover, MSCs reside in a hypoxic microenvironment that can promote apoptosis which leads to therapeutic potential under in vivo conditions (16) by secreting water-soluble immunomodulatory factors (17–19).

There is an urgent need to augment the efficacy of MSCs-based cellular therapy by tinkering apoptosis in conjunction with hypoxia appears to be the most effective conditions that impact immunomodulation potential, particularly for GVHD patients who do not respond favorably to viable MSCs. In view of this prerequisite, our study provides experimental evidence that apoptotic in conjunction with hypoxia enhances the immune regulatory potential of tissue-specific human MSCs (BM, WJ) in a cellular model of aGVHD.

## MATERIAL AND METHODS

### Ethics approval

The study involving mesenchymal stem cells from human was approved by both the Institutional Ethics Committee (IECPG-542/23.09.2020) and the Institutional Committee for Stem Cell Research and Therapy (IC-SCR/110/20(R). Informed written consents were obtained from all participating subjects and all procedures were performed as per the guidelines and regulations approved by the ethics committee.

### Isolation of MSCs from human Bone Marrow and Wharton’s Jelly

Human BM aspirates were collected from healthy donors (n=5) of allogeneic stem cell transplant recipients at the Department of Medical Oncology, Dr. B. R. Ambedkar Institute Rotary Cancer Hospital, AIIMS, New Delhi, and MSCs were isolated using our previously established protocol (20,21). Briefly, the aspirate was seeded in a 60mm culture dish (Nunc^TM^, Thermo Fisher Scientific, USA) with 1X Low Glucose-Dulbecco’s Modified Eagle Media (LG-DMEM) (Thermo Fisher Scientific, USA) supplemented with 10% FBS (Thermo Fisher Scientific, USA), 1% antibiotic-antimycotic solution (Thermo Fisher Scientific, USA), and incubated in a humidified chamber at 37C with 5% CO_2_, 21% O_2_. After 72 hours, the media was replaced with LG-DMEM complete media with Stem Pro MSC Serum-Free Media (SFM) (Thermo Fisher Scientific, USA) in a 3:1 ratio, and the media was changed every third day until the cell confluency reached 80%. Adherent cells were trypsinized using 0.05% Trypsin-EDTA and reseeded for further experiments.

Human umbilical cord (UC) (n=5) was collected in a sterile transport media from the Department of Obstetrics and Gynaecology, AIIMS, New Delhi, and MSCs were isolated using our previously established protocol (20). Briefly, the explant culture method was used to obtain MSCs from Wharton’s Jelly (WJ) and approximately 2cm*2cm size were seeded in a 35mm culture dish (Nunc^TM^, Thermo Fisher Scientific, USA) and the coverslips were placed over explants and 1X LG-DMEM complete medium was added dropwise. The culture dish was placed in a humidified chamber at 37C, 5% CO_2_ for 2-3 hours for the attachment of explants followed by the addition of 1X LG-DMEM with Stem Pro MSC SFM in a 3:1 ratio. The cells were harvested using 0.05% Trypsin-EDTA after the attainment of 80% cell confluency and the cells were expanded for further experiments.

### Characterization of tissue-specific MSCs

Both BM-MSCs and WJ-MSCs (Passage-3) were characterized according to the International Society for Cellular Therapy (ISCT) guidelines (22) using our established protocols (23). Briefly, MSCs were characterized for their plastic adherence, surface marker profile, and trilineage differentiation potential. Further, the metabolic rate of MSCs was determined on Days 1, 3, 5, and 7 using an MTT assay, and the population doubling time (PDT) was enumerated after 72 hours using the following formula (24):

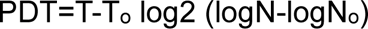

where, T: Time of harvesting, T_o_: Time of seeding, N: Number of cells harvested, N_o_: Number of cells seeded

### Generation and characterization of tissue-specific hypoxia-preconditioned apoptotic MSCs

MSCs were pooled to attain homogeneity and passages 3-5 were used for subsequent in vitro experiments. Hypoxia-preconditioned apoptotic MSCs were generated either sequentially or concurrently. In the sequential method, MSCs were first preconditioned with 1% O_2_ for 24 hours, followed by treatment with 1µM staurosporine (STS) (Sigma, USA) for 24 hours in 1X LG-DMEM complete media in a tri-gas incubator (Thermo Fisher Scientific, USA), termed as MSCs^HYP→APO^. In the concurrent method, MSCs were exposed to 1% O_2_ and 1µM STS for 24 hours in a tri-gas incubator, termed MSCs^HYP+APO^. Additionally, MSCs were exposed to 1% O_2_ alone for 24 hours, termed hypoxia preconditioned MSCs (MSCs^HYP^).

The respective cell suspensions were collected, washed, and stained with Annexin V/7AAD for the enumeration of apoptotic cells using a DxFlex flow cytometer (Beckmann Coulter), and the data was analyzed using Kaluza software version 2.1 (Beckmann Coulter). Additionally, the apoptosis was confirmed by cleaved caspase-3 (Cell Science Technology, USA) using western blotting (10).

### T-cell proliferation assay

Peripheral blood (PB) was collected from patients with grade II-IV acute graft-versus-host-disease (aGvHD) (n=5) in a sterile sodium heparin-coated vacutainer (BD Biosciences, US) and the peripheral blood mononuclear cells (PBMNCs) were isolated using a standardized protocol (25). Briefly, PB was diluted with 1X PBS in a 1:1 ratio, followed by layering of diluted blood over Histopaque-1077 density gradient medium (Sigma, USA) in a 2:1 ratio and centrifuged at 400g for 30 minutes at room temperature. The buffy coat containing mononuclear cells was collected and washed with 1X PBS (Thermo Fisher Scientific, USA). CD3^+^ T-cell was isolated from PBMNCs by negative selection using a Pan T cell isolation kit, human (Miltenyi Biotec, USA) as per the manufacturer’s instructions. CD3^+^ T-cell was labeled with 1µM Cell Trace^TM^ CFSE dye (BD Biosciences, USA) for 20 minutes at 37C and followed by activation of CD3^+^ T-cell with PHA (1µg/ml) (Sigma, USA) and IL-2 (50IU/ml) (Thermo Fisher Scientific, USA) in 1X Rosewell Park’s Memorial Institute (RPMI) medium supplemented with L-glutamine (Thermo Fisher Scientific, USA), 10% FBS, 1% antibiotic-antimycotic solution at 37C, 5% CO_2_ for 48 hours.

For co-culture, hypoxia-preconditioned MSCs (MSCs^HYP^, MSCs^HYP→APO^, MSCs^HYP+APO^) were treated with mitomycin-C (15µg/ml) (Thermo Fisher Scientific, USA) for 1 hour in 1X LG-DMEM incomplete medium. The mitomycin-treated MSCs were co-cultured with CFSE-labeled activated T-cell in a 1:10 ratio in 1X RPMI complete medium with 1X LG-DMEM complete medium in a 1:1 ratio for 3 days. After 3 days of co-culture, cells were collected, washed with 1X PBS, and stained with fluorochrome-conjugated anti-human monoclonal antibodies for CD3, CD4, CD8, and CD45 (Beckmann Coulter) for 30 minutes at room temperature and the proliferation of T-cell was assessed using a DxFlex flow cytometer (Beckmann Coulter, USA) and the data was analyzed using Kaluza software version 2.1 (Beckmann Coulter, USA). CFSE-labeled activated T-cell in the absence of MSCs was used as a control for the normalization of the proliferation of T-cell in the direct co-culture (26,27).

### Induction of Tregs

The mitomycin-treated MSCs and activated T-cell were co-cultured in a 1:10 ratio for 5 days. After 5 days of co-culture, cells were collected, washed with 1X PBS, and stained with fluorochrome-conjugated anti-human monoclonal antibodies against CD3, CD4, CD8, CD25, CD45 (Beckmann Coulter, USA). For intracellular staining, cells were fixed and permeabilized with a FOXP3 buffer set (BD Biosciences, USA) followed by staining with anti-FOXP3 monoclonal antibody for 30 minutes at room temperature. A minimum of 50,000 events were acquired by a DxFlex flow cytometer (Beckmann Coulter) and the data was analyzed using Kaluza software version 2.1 (Beckmann Coulter). Activated T-cell in the absence of MSCs was used as a control for the baseline expression of CD3^+^ CD4^+^ CD25^+^ FOXP3^+^ Tregs (28,29).

### Enumeration of effector memory helper T cell subtypes (Th1, Th2, and Th17)

The proportion of Th1, Th2, and Th17 were enumerated in the 5-day co-culture of mitomycin-treated MSCs and activated T-cell using flow cytometry. The cells were initially stained with fluorochrome-conjugated antihuman monoclonal antibodies targeting chemokine receptors (CXCR3, CCR4, CCR7, CCR6) at 37C for 30 minutes followed by staining anti-human CD3, CD4, CD8, CD45RA, CD45 monoclonal antibodies and Th1, Th2, Th17 was enumerated using DxFlex flow cytometer (Beckmann Coulter) (28,30).

### Macrophage polarization

CD14^+^ monocytes were isolated from PBMNCs using a pan-monocyte isolation kit, human (Miltenyi Biotec, USA) following the manufacturer’s guidelines. The isolated monocytes were treated with 25ng/ml GM-CSF for 5 days to induce differentiation into M0 macrophages. Subsequently, M0 macrophages were treated with 10ng/ml LPS and 50ng/ml IFN-γ for 24 hours to generate M1 macrophages. The mitomycin-treated MSCs were co-cultured with M1 macrophages in a 1:10 ratio for 3 days. The polarization of macrophages from M1 to M2 phenotype was assessed by staining the cells with fluorochrome-conjugated anti-CD206 (BD Biosciences, USA) followed by fixation and permeabilization of the cells using an Intra Prep Permeabilization kit (Beckmann Coulter, USA) for intracellular staining. Further, the cells were stained with iNOS (Thermo Fisher Scientific, USA), CD68, and Arginase-1 (BD Biosciences, USA) for 30 minutes at room temperature and the cells were acquired using a DxFlex flow cytometer (Beckmann Coulter) to enumerate the M1 and M2 macrophages (31).

### In vitro MSCs clearance assay

CFSE labeled aGvHD patients-derived M1 macrophages were treated with pH Rhodo red succinimidyl ester labeled MSCs in a 1:2 ratio for 24 hours. The percentage of M1 MФ^CFSE+^ was also positive for pH Rhodo red signal used for the calculation of uptake of MSCs by M1 macrophages using a DxFlex flow cytometer (Beckmann Coulter) (9,32).

### Gene expression analysis of immunomodulatory molecules

Total RNA was extracted from the co-culture of MSCs and aGvHD patients derived PBMNCs using the phenol-chloroform extraction method by TRIzol^TM^ reagent (Thermo Fisher Scientific, USA) according to the manufacturer’s protocol. 2 μg total RNA was reverse transcribed to give cDNA using the High-Capacity cDNA Reverse Transcription Kit (Applied Biosystems, Invitrogen, USA). Quantitative real-time polymerase chain reaction (qRT-PCR) was performed using a CFX96 Real-Time System (Bio-Rad). Glyceraldehyde-3 phosphate dehydrogenase (GAPDH) was used as the housekeeping gene to normalize the gene expression. qRT-PCR was performed in triplicate using SYBR Green Master Mix (Promega, US) according to the manufacturer’s instructions. The comparative 2−ΔΔCt method was performed to evaluate the mRNA expression of IDO, PGE2, IL-10, IFN-γ, IL-6, TNF-α, IL-12β, IL-1β and the sequence of primers were listed in Table S1.

### Statistical analysis

All statistical analyses were conducted using GraphPad Prism version 8.4.3. One-way and Tukey’s post hoc tests were used to compare three or more groups. Data was shown as Mean±S.D. and a p-value of ≤ 0.05 was considered statistically significant.

## RESULTS

### Hypoxia preconditioning maintained characteristics of MSCs

Initially, we characterized naïve and hypoxia-preconditioned MSCs (BM, WJ) according to the ISCT guidelines. We observed that hypoxia maintained the parental characteristics of MSCs like plastic adherence, fibroblast-like spindle-shaped morphology, exhibited ≥95% expression of CD105, CD73, CD90, CD29, HLA I, and the ≤2% expression of HLA II, CD34, and CD45, and differentiation into mesodermal lineages (adipogenic, osteogenic, and chondrogenic) (Figure 1A, B, C, S1). While, we did not observe a significant variation in the doubling time of BM-MSCs between normoxia and hypoxia culture conditions (36.21±0.852, 35.93±0.226), however, WJ-MSCs exhibited a significant reduction in doubling time under hypoxia compared to normoxia conditions (22.123±0.227 vs. 26.16±0.284; p ≤ 0.0001) (Figure 1D). Moreover, we observed that both MSCs exhibited exponential growth from days 1 to 5, followed by a decline in proliferation from day 5 to day 7 under normoxia and hypoxia culture conditions (Figure 1E). Overall, WJ-MSCs demonstrated higher proliferation rates than BM-MSCs under both normoxia and hypoxia conditions, as evidenced by both PDT and metabolic rate. Low oxygen level during hypoxia is sensed by the Von Landau Hipple factor and manifested by the upregulation of the HIF-1α transcription factor which orchestrates hypoxia-induced changes in the cells. HIF-1α gene which undergoes ubiquitination in normoxic conditions but becomes stabilized following exposure to hypoxia. Consequently, the expression levels of this gene serve as an indicator of the cellular response to hypoxic conditions (33). Interestingly, both MSCs (BM, WJ) had higher expression of HIF-1α from 6 hours (5.085±0.0495, 8.349±0.454-fold change) to 12 hours (9.65±0.161, 14.420±0.665 fold change) followed by a decline in their expression from 24 hours (6.382±0.059, 10.324±0.119 fold change) to 48 hours (4.288±0.656, 5.470±0.334 fold change) at the gene and protein level (Figure 1F, G).

**Figure 1:**
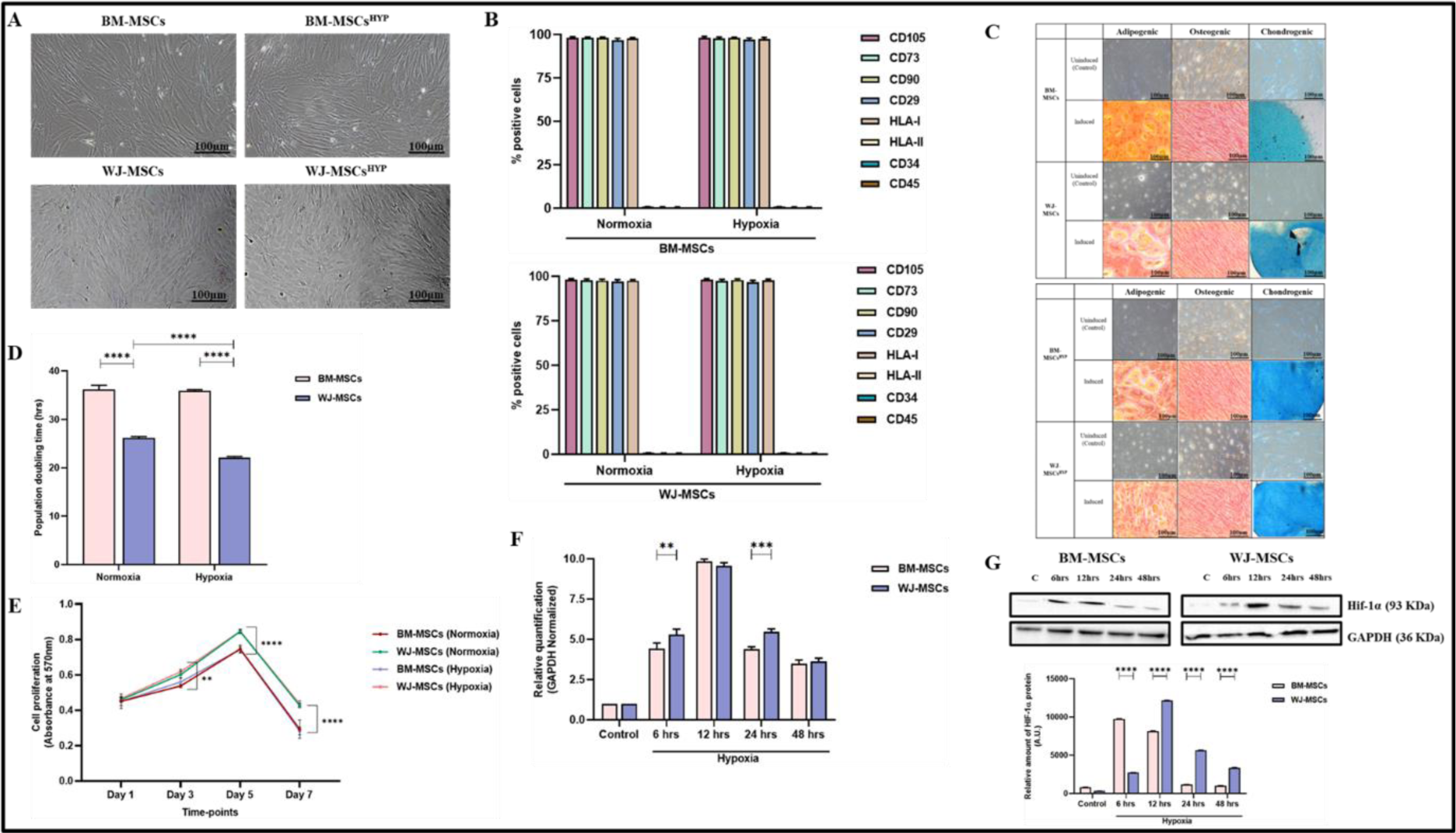
Characterization of human MSCs (BM-MSCs, BM-MSCs^HYP^, WJ-MSCs, and WJ-MSCs^HYP^) (A) Morphological images. (B) A bar graph depicts surface marker profiling using flow cytometry. (C) Trilineage differentiation into adipocytes (28 days of induction), osteocytes (21 days of induction), and chondrocytes (14 days of induction). (D) The bar graph represents the population doubling time in hours. (E) The line diagram represents the growth kinetics. (F) The bar graph depicts the relative expression of the HIF-1α gene using qPCR before and after exposure to 1% O_2_ at various time intervals (6h, 12h, 24h, 48h). (G) Western blot images represent the expression of Hif-1α protein and the bar graph represents the relative expression of Hif-1α protein before and after exposure to 1% O_2_ at various time intervals (6h, 12h, 24h, 48h). Data shown as Mean±S.D.; Statistical analysis: Tukey’s multiple comparisons test; ***≤0.001; ****≤0.0001. All experiments were done in triplicates. Scale bar: 10X=100µm. *Abbreviations: BM: Bone marrow; WJ: Wharton’s Jelly; MSCs: Mesenchymal stem cells; HYP: Hypoxia-preconditioned*.

Recent findings challenge the significance of cell viability for immunomodulation potential, though, MSCs undergo apoptosis post-administration within the host (6,8,34), and hypoxia augments the apoptosis of MSCs (17). Therefore, we induced apoptosis in hypoxia-preconditioned MSCs (BM, WJ) using STS in both successive and simultaneous ways. Both approaches resulted in a notable alteration in the morphology of MSCs, characterized by cellular shrinkage, blebbing, and fragmentation (Figure 2A) with the proportion of apoptotic MSCs (≥98%), confirmed by Annexin-V/7AAD staining (Figure 2B, S2) and the expression of cleaved caspase-3 was enhanced in apoptotic MSCs (Figure 2C). Interestingly, despite apoptosis, the phenotype of MSCs in both approaches, remained unchanged (Figure 2D, S3) indicating the stability of attributes of hypoxia-preconditioned MSCs following apoptosis.

**Figure 2:**
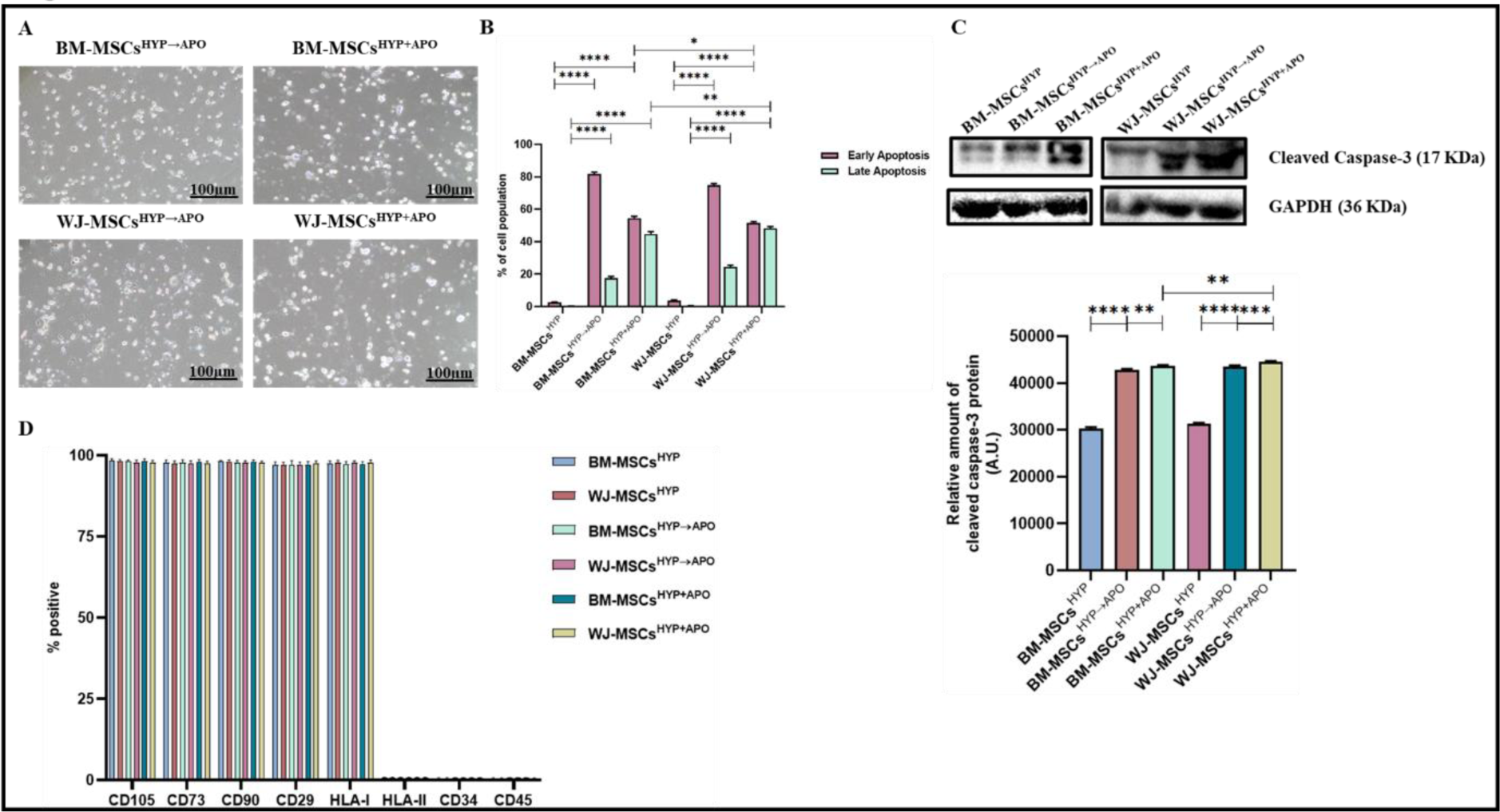
Characterization of apoptotic human MSCs (BM-MSCs^HYP→Apo^, BM-MSCs^HYP+APO^, WJ-MSCs^HYP→APO^, and WJ-MSCs^HYP+APO^. A) Microscopic images. B) A bar graph represents the percentage of apoptotic cells (early and late apoptotic cells). C) Western blot images depicts expression of cleaved caspase-3 and bar grapgh represents the relative expression of cleaved caspase-3. D) The bar graph represents surface expression profile using flow cytometry. Data shown as Mean±S.D.; Statistical analysis: Tukey’s multiple comparisons test; ****≤0.0001. All experiments were done in triplicates. Scale bar: 10X=100µm. *Abbreviations: BM: Bone marrow; WJ: Wharton’s Jelly; MSCs: Mesenchymal stem cells; APO: Apoptosis; HYP: Hypoxia-preconditioned*.

### WJ-MSCs^HYP+APO^ induced immune programming of effector T-cell

MSCs not only manifest regenerative potential but also play a crucial role in immunomodulation, contributing significantly to the maintenance of immune homeostasis and preventing inflammation in the host. To address this, we assessed whether hypoxia-preconditioned both viable and apoptotic MSCs (MSCs^HYP^, MSCs^HYP→APO^, MSCs^HYP+APO^) would be able to polarize effector T-cell and macrophages or not. Initially, we co-cultured CD3^+^ T-cell with MSCs and T-cell proliferation, induction of Tregs, and polarization of helper T-cell from pro-inflammatory to anti-inflammatory phenotype was assessed using flow cytometry. We observed that both MSCs^HYP^ (BM, WJ) inhibited the proliferation of T-cell significantly, wherein, WJ-MSCs^HYP^ were more effective in suppressing T-cell proliferation than BM-MSCs^HYP^ (65.62% vs. 73.964%; p ≤0.0001). Interestingly, hypoxia-preconditioned apoptotic MSCs (MSCs^HYP→APO^, MSCs^HYP+APO^) did not effectively inhibit T-cell proliferation, regardless of the tissue source (Figure 3A, S4). Similarly, MSCs^HYP^ (BM, WJ) increased CD4/CD8 ratio indicating the inhibitory impact of MSCs on CD8^+^ T-cell populations (Figure 3B).

**Figure 3:**
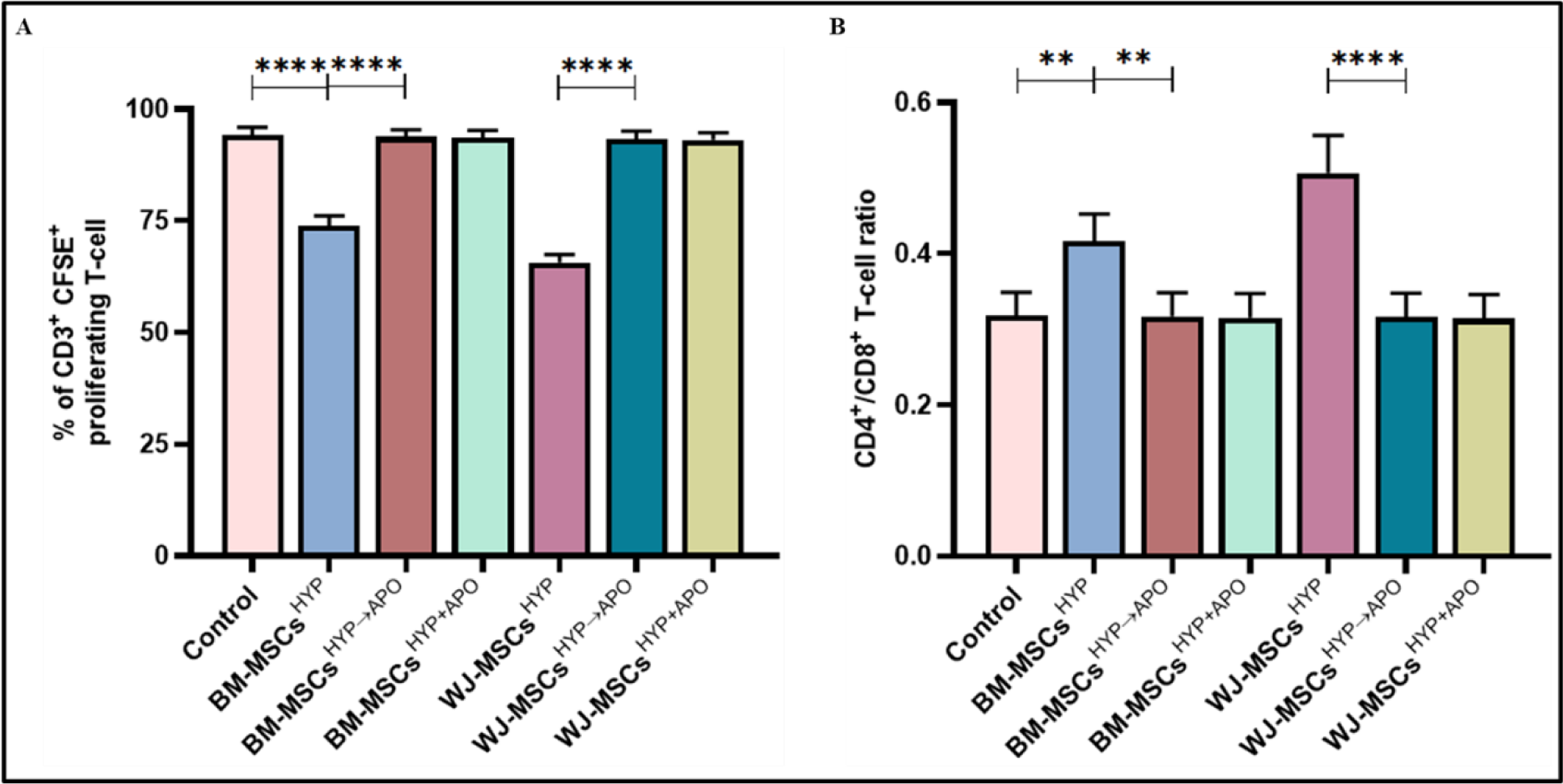
Effect of hypoxia-preconditioned MSCs (MSCs^HYP^, MSCs^HYP→APO^, MSCs^HYP+APO^) on the proliferation of aGVHD patients derived T-cell. The bar graph represents (A) the percentage of CD3^+^ CFSE^+^ proliferating T-cell (n=5). (B) the ratio of CD4^+^/CD8^+^ T cell (n=5) in the direct co-culture of MSCs and T-cell. Data shown as Mean±S.D.; Statistical analysis: Tukey’s multiple comparisons test; **≤0.01; ****≤0.0001. *Abbreviations: BM: Bone marrow; WJ: Wharton’s Jelly; MSCs: Mesenchymal Stem Cells; HYP: Hypoxia-preconditioned; APO: Apoptosis*.

Based on the MSC^HYP^-mediated loss of the CD8^+^ T-cell population and concomitant increase in CD4^+^ T-cell, we anticipated that MSCs might favor either proliferation of CD4^+^ T-cell or promote their differentiation into Tregs as one of the underlying immune polarization mechanisms. To demonstrate this, we also analyzed the Tregs population in the direct co-culture, as described above. Following our expectation, the co-culture of MSCs^HYP^ (BM, WJ) enhanced the Treg populations wherein WJ-MSCs^HYP^ enhanced the differentiation of CD4+ T-cell to Tregs than BM-MSCs^HYP^ (10.08% vs. 5.856%; p: ≤0.0001). In contrast and unlike T cell proliferation, hypoxia-preconditioned apoptotic MSCs (MSCs^HYP→APO^, MSCs^HYP+APO^) enhanced the differentiation of CD4^+^ T-cell into Tregs synergistically over MSCs^HYP^, regardless of tissue source. However, the percentage of Tregs induced by WJ-MSCs^HYP+APO^ was significantly higher than WJ-MSCs^HYP→APO^ (16.988% vs. 10.954%; p ≤0.0001), with a similar trend observed in BM-MSCs (12.442% vs. 7.929%; p ≤0.0001) (Figure 4A, S5), signify simultaneous exposure of hypoxia and apoptosis to MSCs surpassed successive approach (Figure 4A, S5). These results, supporting our hypothesis, suggested that MSCs are effective in modulating T-cell phenotypically in co-cultures.

**Figure 4:**
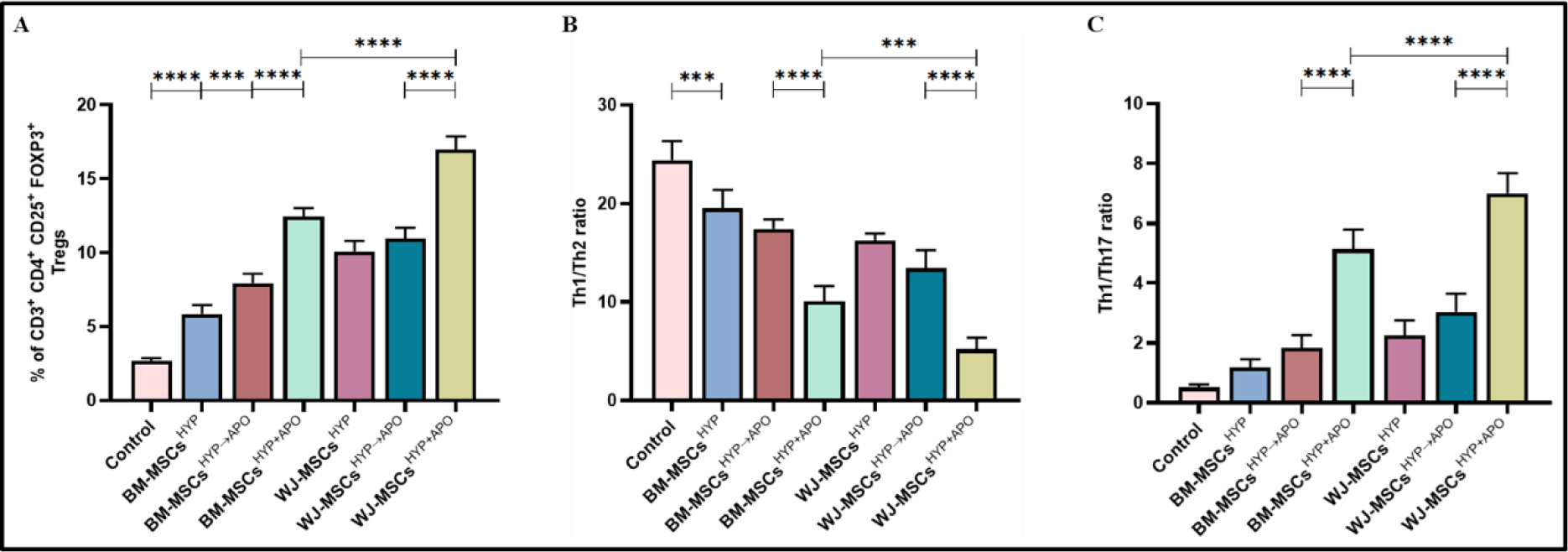
Effect of hypoxia-preconditioned MSCs (MSCs^HYP^, MSCs^HYP→APO^, MSCs^HYP+APO^) on the induction of Tregs and polarization of helper T cell from pro-inflammatory (Th1, Th17) to anti-inflammatory (Th2) phenotype. The bar graph represents (A) the induction of CD3^+^ CD4^+^ CD25^+^ FOXP3^+^ Tregs (n=5). (B) the ratio of Th1/Th2 (n=5). (C) Th1/Th17 ratio (n=5) in the direct co-culture of MSCs and T-cell. Data shown as Mean±S.D.; Statistical analysis: Tukey’s multiple comparisons test; ***≤0.001; ****≤0.0001. *Abbreviations: BM: Bone marrow; WJ: Wharton’s Jelly; MSCs: Mesenchymal stem cells; HYP: Hypoxia-preconditioned; APO: Apoptosis, Tregs: Regulatory helper T-cell, Th: Helper T cell*.

Based on this, we anticipated that these MSCs might tweak them functionally as well. To substantiate this, we analyzed both Th1/Th2 and Th1/Th17 ratios as indicators of overall immune responses. In line with this and following our hypothesis, co-culture of MSCs^HYP^/MSCs^HYP→APO^/MSCs^HYP+APO^ with T-cell reduced the Th1/Th2 ratio and enhanced the Th1/Th17 ratio compared to control, suggesting that these MSCs can polarize Th1 to Th2 and inhibit Th17 response in the direct co-culture system. Similarly, we observed that both BM-MSCs^HYP+APO^ and WJ-MSCs^HYP+APO^ (10.056 vs. 5.214; p 0.0004) caused a significant decrease in Th1/Th2 ratio than sequentially generated counterparts (17.438 vs 13.416; p 0.0038) (Figure 4B, S6). Additionally, a significant increase in the Th1/Th17 ratio was observed with BM-MSCs^HYP+APO^ and WJ-MSCs^HYP+APO^ (7.008 vs. 5.132; p ≤0.0001) compared to their sequentially generated apoptotic MSCs^HYP→APO^ (3.04 vs.1.836; p ≤0.0001), indicating a shift towards an anti-inflammatory state (Figure 4C, S6).

These results prudently advocated that both MSCs^HYP^ and MSCs ^HYP→APO^/MSCs^HYP+APO^ are capable of T-cell programming which indicated one of the potential mechanisms by which these cells can manipulate host immune responses.

### WJ-MSCs^HYP+APO^ undergo efferocytosis more efficiently and trigger macrophage polarization with a concurrent increase in the expression of immunomodulatory molecules

Indeed, T cells are major determinants of effective immune response but these in turn need innate immune cells which are capable of deciding the fate of immune responses. Mainly, macrophages play a major role in organ transplant where these cells scavenge and clear dead cells from tissue for homeostasis. To address this, we assessed the phagocytosis of MSCs^HYP^, MSCs^HYP→APO^, and MSCs^HYP+APO^ by macrophages by a process known as efferocytosis under *in vitro* settings. To mimic this in our model, we co-cultured MSCs with macrophages, as described above, and allowed them to phagocytize and correlate efferocytosis using flow cytometry. In line with the hypothesis, our data revealed that macrophages exhibited enhanced efficiency of efferocytosis of both MSCs^HYP→APO^ and MSCs^HYP+APO^ over MSCs^HYP^, irrespective of tissue origin. Our results demonstrated that BM-MSCs^(HYP+APO)^ and WJ-MSCs^(HYP+APO)^ (86.588% vs. 95.262%; p ≤0.0001) than BM-MSCs^(HYP→APO)^ and WJ-MSCs^(HYP→APO)^ (64.53% vs. 76.062%; p ≤0.0001), thereby explaining the maximum immunomodulation mediated by WJ-MSCs^(HYP+APO)^ (Figure 5A, S7). Frequent apoptosis and uptake of various bacteria/dead cells carrying neutrophils and/or fibroblast and other cells by tissue-resident scavenging and CD68^+^ effector macrophages trigger the release of TGF-β, prostaglandins and other mediators which further promote in situ polarization of effector immune cells toward their refractory/anti-inflammatory phenotype. Since we have seen Th2 bias in MSCs^HYP+APO^/MSCs^HYP→APO^ co-cultured T cells and increased efferocytosis of MSCs^HYP+APO^/MSCs^HYP→APO^ by macrophages, we anticipated a potent in situ programming of M1 effector macrophages to their M2 phenotype. To address this, we analyzed the M1/ M2 population in the macrophages which were co-cultured with MSCs^HYP^/MSCs^HYP+APO^/MSCs^HYP→APO,^ and a panel of Th1/Th2 effectors in the direct co-culture of MSCs^HYP^/MSCs^HYP+APO^/MSCs^HYP→APO^ with CD3^+^ T-cell.

**Figure 5:**
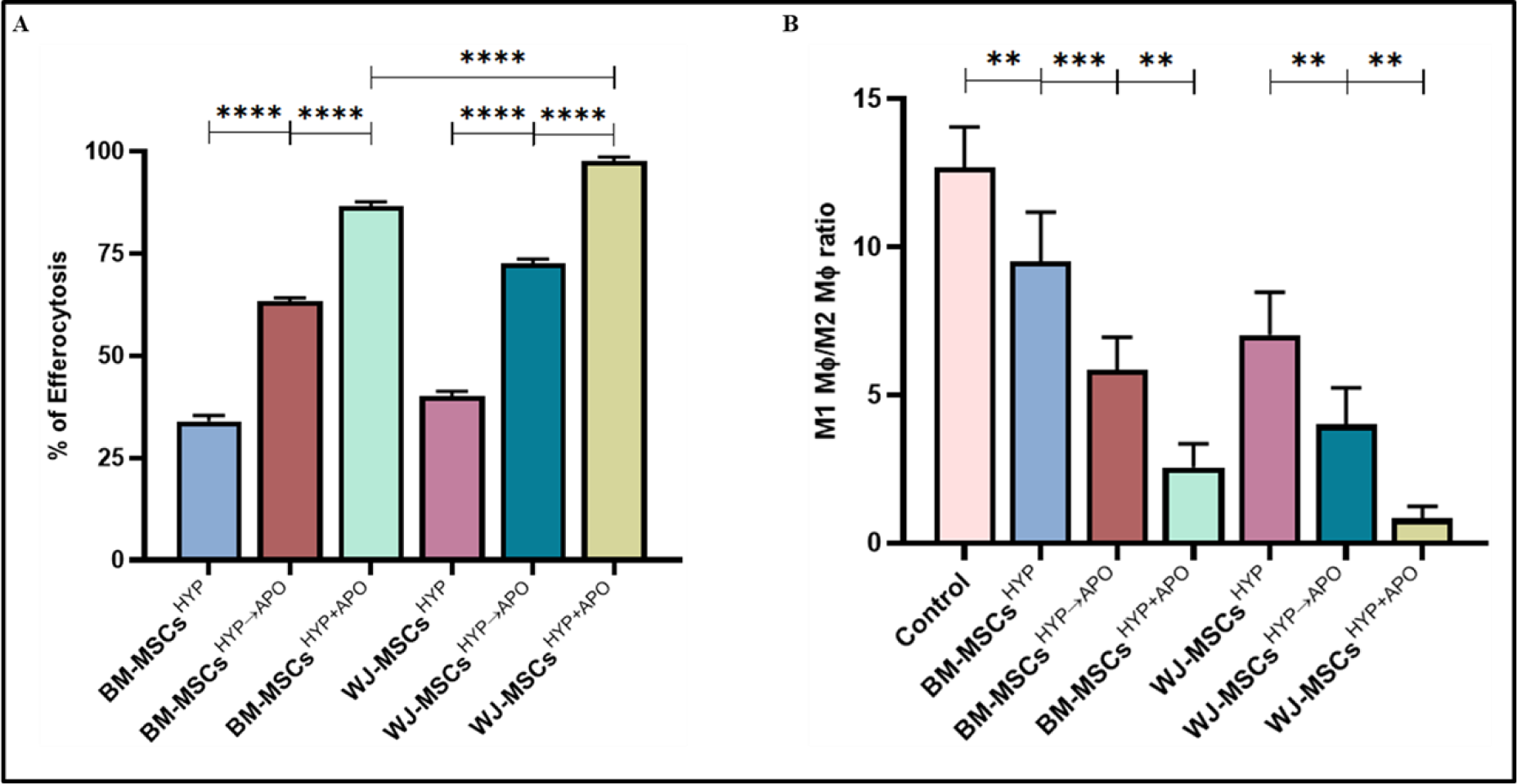
Effect of hypoxia-preconditioned MSCs (MSCs^HYP^, MSCs^HYP→APO^, MSCs^HYP+APO^) on the efferocytosis of MSCs and macrophage polarization from proinflammatory (M1) to anti-inflammatory (M2) phenotype. The bar graph represents (A) efferocytosis of MSCs by macrophages (n=5). (B) the ratio of M1 MФ/M2 MФ in the co-culture of MSCs and M1 MФ (n=5). Data shown as Mean±S.D.; Statistical analysis: Tukey’s multiple comparisons test; *≤0.05; **≤0.01; ***≤0.001; ****≤0.0001. *Abbreviations: BM: Bone marrow; WJ: Wharton’s Jelly; MSCs: Mesenchymal stem cells; HYP: Hypoxia-preconditioned; APO: Apoptosis; MФ: Macrophages*.

Following our hypothesis, our data prudently demonstrated a significant polarization of the M1 (iNOS^+^ Arginase-1^-^) macrophages toward the M2 (iNOS^-^ Arginase-1^+^) phenotype with MSCs^(HYP+APO)^ compared to MSCs^(HYP→APO)^ in both BM (2.860 vs. 5.857; p 0.0031) and WJ (0.817 vs. 4.004; p 0.0045), evident by a decrease in the M1/M2 ratio (Figure 5B, S8). Furthermore, our experiments provided experimental evidence that co-culture of WJ-MSCs^HYP+APO^ enhanced the expression of immunomodulatory molecules which are capable of opposing Th1 effector cytokines (IDO, PGE2), inhibited the expression of Th1 effector (IL-1β, IL-12β, TNF-α, IL-6), and enhanced the expression of Th2 effector cytokines (IL-10) providing evidence of immune metabolic programming of effectors immune cells. (Figure 6A-H).

**Figure 6:**
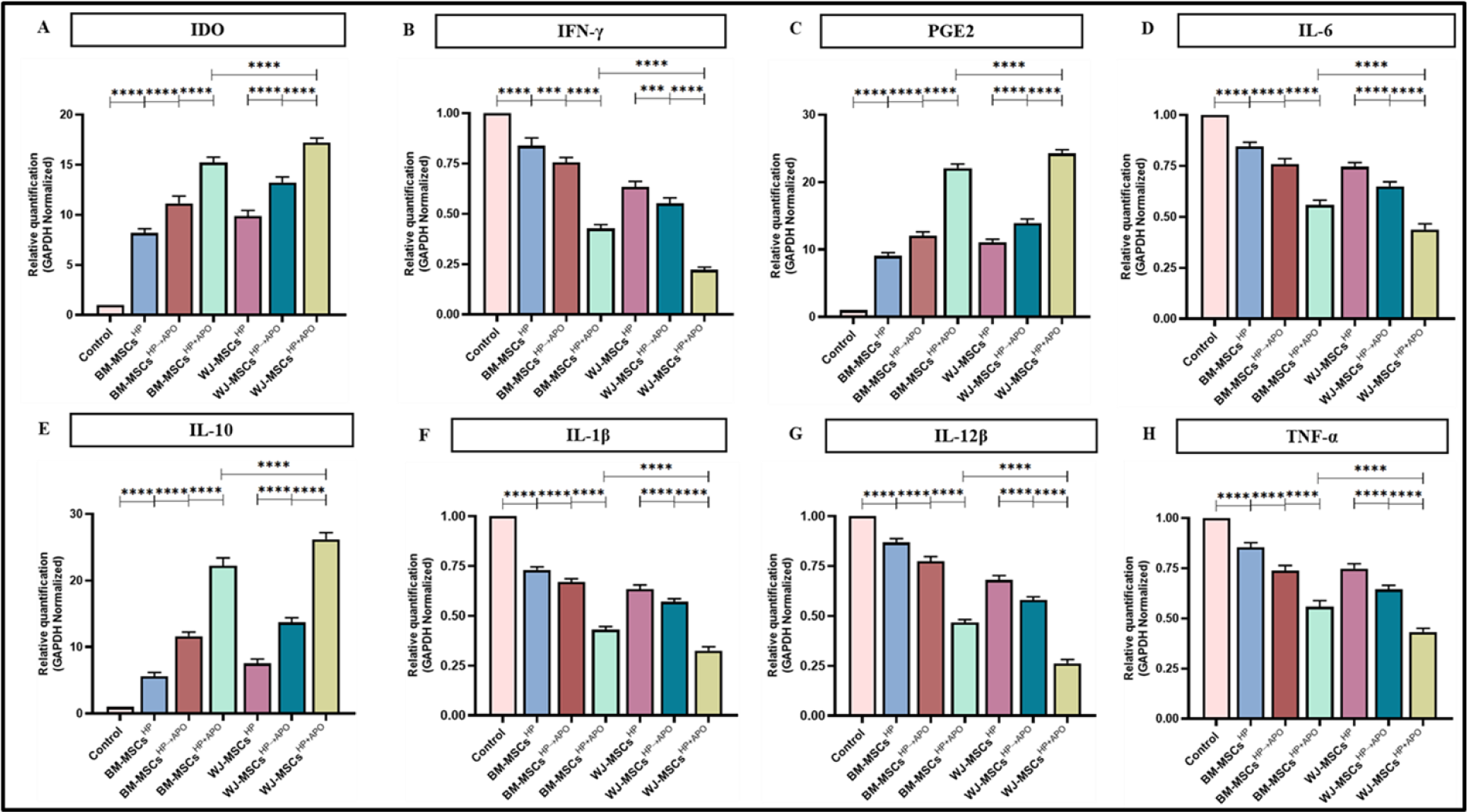
Effect of hypoxia-preconditioned MSCs (MSCs^HYP^, MSCs^HYP→APO^, MSCs^HYP+APO^) on the relative expression of immunomodulatory molecules and cytokines. The bar graphs represent relative mRNA expression of (A) IDO (B) IFN-γ (C) PGE2 (D) IL-6 (E) IL-10 (F) IL-1β (G) IL-12β (H) TNF-α in the direct co-culture of MSCs and aPBMNCs derived from aGVHD patients (n=5). Data shown as Mean±S.D.; Statistical analysis: Tukey’s multiple comparisons test; *≤0.05; **≤0.01; ***≤0.001; ****≤0.0001. *Abbreviations: BM: Bone marrow; WJ: Wharton’s Jelly; MSCs: Mesenchymal stem cells; HYP: Hypoxia-preconditioned; APO: Apoptosis; IDO: Indoleamine 2,3 dioxygenase; PGE2: Prostaglandin E2; IFN-γ: Interferon-γ; IL: Interleukin; TNF-α: Tumor necrosis factor-α*.

## DISCUSSION

Recent studies have suggested that instead of undergoing in situ de/re-differentiation, transplanted MSCs undergo apoptosis following administration, primarily as a result of interactions with the host’s immune effector cells and ultimately leading to immune modulation of the host (6,36). Thus, maintaining the parental identity of MSCs while enhancing their immunomodulatory potential is vital, highlighting the importance of modifying MSCs before their therapeutic application. In this context, our study shed new light on how the preconditioning of MSCs with short-term hypoxia (1% O_2_) and inducing apoptosis can enhance the immunomodulatory potential of MSCs (BM, WJ). Our results indicated that preconditioning of MSCs with hypoxia (1% O_2_) for 24 hours did not affect the fibroblast-like spindle-shaped morphology of MSCs, which is consistent with the previous findings (37). However, several studies demonstrated that hypoxia maintained the expression of negative markers (CD34, CD45, HLA-II) with a variable impact on the expression of positive markers (CD105, CD73, CD90, CD29, HLA-I) (38–42). In contrast, few studies reported that a hypoxic environment did not confer a significant change in surface profile (43). In this line, we also observed that a short exposure of MSC to hypoxia did not cause any change in surface markers on MSCs. Multiple studies revealed that hypoxia has a negative impact on the clonogenic potential of MSCs (WJ, adipose tissue) (43,44). However, our observations revealed differences in PDT with no variation in BM-MSCs while a significant difference was observed in WJ-MSCs under both normoxia and hypoxia culture conditions, consistent with previous findings (33,45). The observed variations are probably a result of the diverse protocols employed, variations in culture media compositions, oxygen levels, and the inherent heterogeneity among donors (40). Notably, there was no difference in the differentiation of MSCs into mesodermal lineages (adipogenic, osteogenic, and chondrogenic) across both culture conditions in contrast to previous studies which demonstrated that reduced oxygen pressure decreased the differentiation of MSCs into adipogenic, osteogenic with an increase in their chondrogenic lineage (46,47). Interestingly, MSCs^HYP^ had higher expression of HIF-1α from 6 to 12 hours followed by a decline in their expression from 24 to 48 hours at the gene and protein level consistent with our previous findings (33). We observed a significant alteration in the morphology of hypoxia-preconditioned apoptotic MSCs similar to the previous observations (10,11,48,49).

Several studies reported that hypoxia-preconditioned MSCs had a profound effect on the suppression of T-cell proliferation, induction of CD4^+^ or CD8^+^ regulatory T-cell, and modified T-cell polarization toward the anti-inflammatory milieu in inflammatory conditions like GVHD and also by increasing the apoptosis of MSCs (42,50–61). Similarly, we observed that WJ-MSCs^HYP^ exhibited greater efficiency in inhibiting aGvHD patients-derived T-cell proliferation than BM-MSCs^HYP^. Apoptotic MSCs exerted immunosuppressive effects in allergic asthma similar to live MSCs (14). In contrast, various preclinical studies have demonstrated that apoptotic MSCs are more potent in promoting immune suppression (8), and it was elucidated that the mode in which apoptotic MSCs are administered significantly influences effector cell infiltration in aGVHD murine model (6). Their findings highlighted that intraperitoneal administration resulted in a more pronounced decrease in effector cell infiltration compared to the intravenous route and apoptotic BM-MSCs did not show a significant inhibition in T cell proliferation under *in vitro* settings (6). Similarly, we observed that MSCs^HYP+APO^/MSCs^HYP→APO^ did not inhibit the proliferation of T-cell regardless of their tissue source while WJ-MSCs^HYP+APO^ exhibited enhanced efficacy in inducing Tregs and this enhanced immunomodulatory potential was further highlighted by their ability to polarize pro-inflammatory T-cell subsets towards an anti-inflammatory phenotype, as evidenced by changes in the Th1/Th2 and Th1/Th17 ratios.

Previous findings revealed that efferocytosis of apoptotic MSCs by macrophages contributes to the programming of macrophages to M2 phenotype with increased secretion of immunomodulatory molecules such as PGE2, IDO, IL-10, PD-L1, and a concurrent decrease in IFN-γ, TNF-α, NO production, resulting in alleviation of inflammation with immune metabolic alterations (8–10,12,14,62–64). We observed that WJ-MSCs^HYP+APO^ demonstrated increased efferocytosis by macrophages and induced polarization towards an M2 phenotype which was associated with upregulated expression of immunomodulatory molecules such as IDO, PGE2, and IL-10, along with a concomitant decrease in pro-inflammatory cytokines, suggesting their role in resolving inflammation and promoting tissue repair.

Nevertheless, despite the emerging evidence challenging the significance of cell viability, it is worth noting that several viable BM-MSCs-based products such as Temcell, and Remestemcel-L have been approved for treating conditions such as aGVHD and steroid-refractory GVHD (SR-GVHD) respectively (65,66). It is observed that only a proportion of patients exhibit a positive response to the therapy. This could suggest that the lack of responsiveness in certain patients might be attributed to inadequate apoptosis of MSCs, potentially stemming from a deficiency in immune cells capable of inducing apoptosis in MSCs (10). To address this issue, our findings highlight that WJ-MSCs^HYP+APO^ is a promising therapeutic strategy for immune-related disorders, and by exploring the immunomodulatory properties of MSCs under hypoxia and apoptosis, it may be possible to enhance their therapeutic efficacy and improve clinical outcomes for patients with conditions such as aGVHD. However, further preclinical and clinical studies are warranted to validate these findings and optimize the protocols for MSCs-based cellular therapy in immune-related diseases.

## AUTHOR’S CONTRIBUTIONS

MM and MM performed the experiments, acquisition, and analysis of data, results interpretation, and wrote the manuscript. SG, SR, RG, and BN were involved in data interpretation and analysis. LK, SB, VD, DP, PSM, RP, MA, AKG, RD, TS, and MM provided patient samples and their clinical details. TDS, SKS, MS, and CPP were involved in data analysis. HP, SM, and RKS conceptualized the study, designed and supervised the experiments, data interpretation and analysis, and wrote the manuscript. All authors critically reviewed, and approved the final version of the manuscript.

## ACKNOWLEDGEMENT

The authors express their gratitude to the All India Institute of Medical Sciences (AIIMS), New Delhi, India for facilitating the execution of the study. A graphical abstract was created using Biorender.com.

---

**Figure S1:**
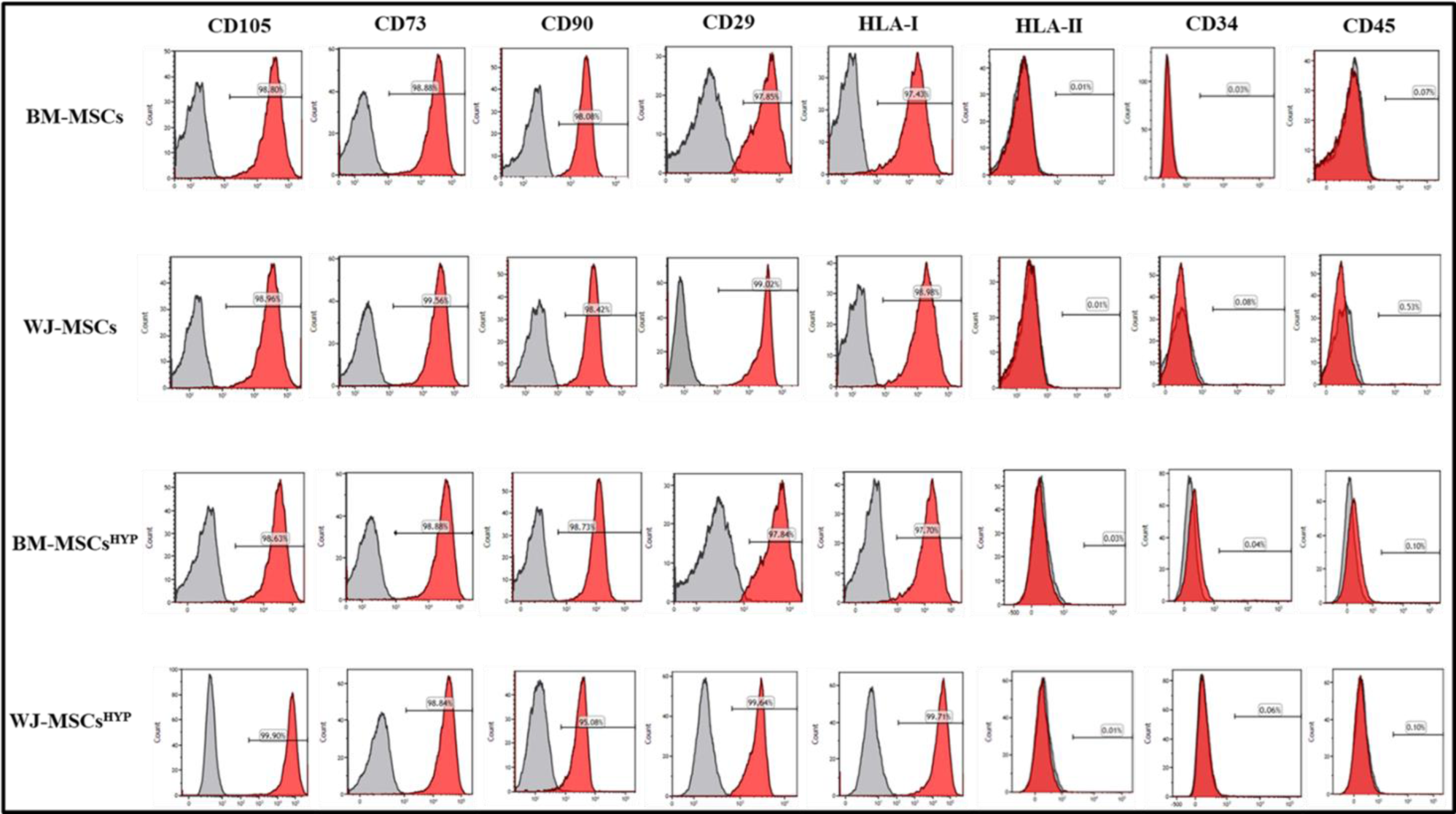
Histograms depicts the surface profile of BM-MSCs, BM-MSCs^HYP^, WJ-MSCs, WJ-MSCs^HYP^ using flow cytometry. *Abbreviations: BM: Bone marrow; WJ: Wharton’s Jelly; MSCs: Mesenchymal Stem Cells; HYP: Hypoxia-preconditioned*.

**Figure S2:**
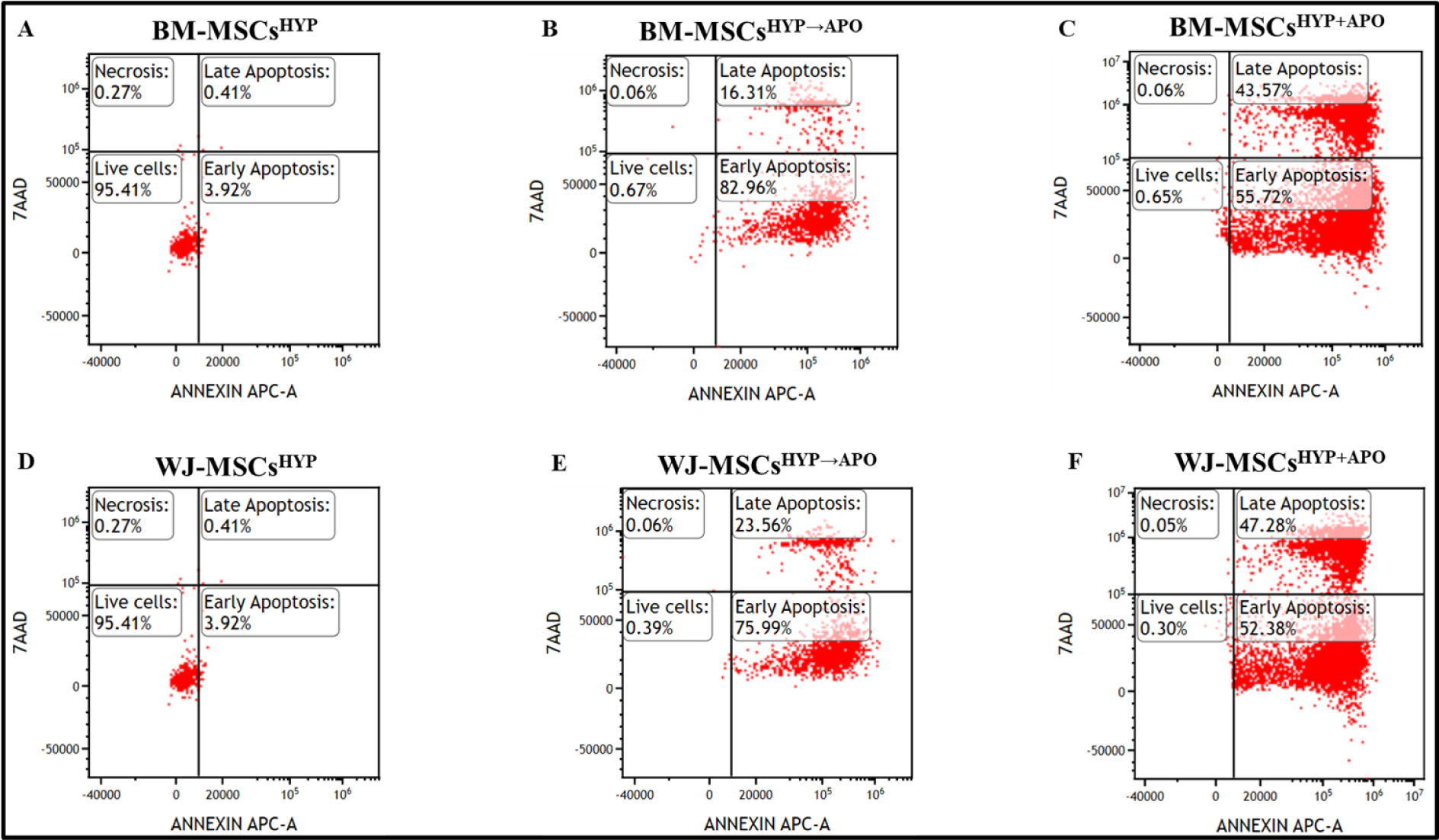
Dot plots depicts the percentage of cell population of (A) BM-MSCs^HYP^. (B) BM-MSCs^HYP→APO^. (C) BM-MSCs^HYP+APO^. (D) WJ-MSCs^HYP^. (E) WJ-MSCs^HYP→APO^. (F) WJ-MSCs^HYP+APO^ using Annexin-V/7AAD staining. *Abbreviations: BM: Bone marrow; WJ: Wharton’s Jelly; MSCs: Mesenchymal Stem Cells; HYP: Hypoxia-preconditioned; APO: Apoptosis*.

**Figure S3:**
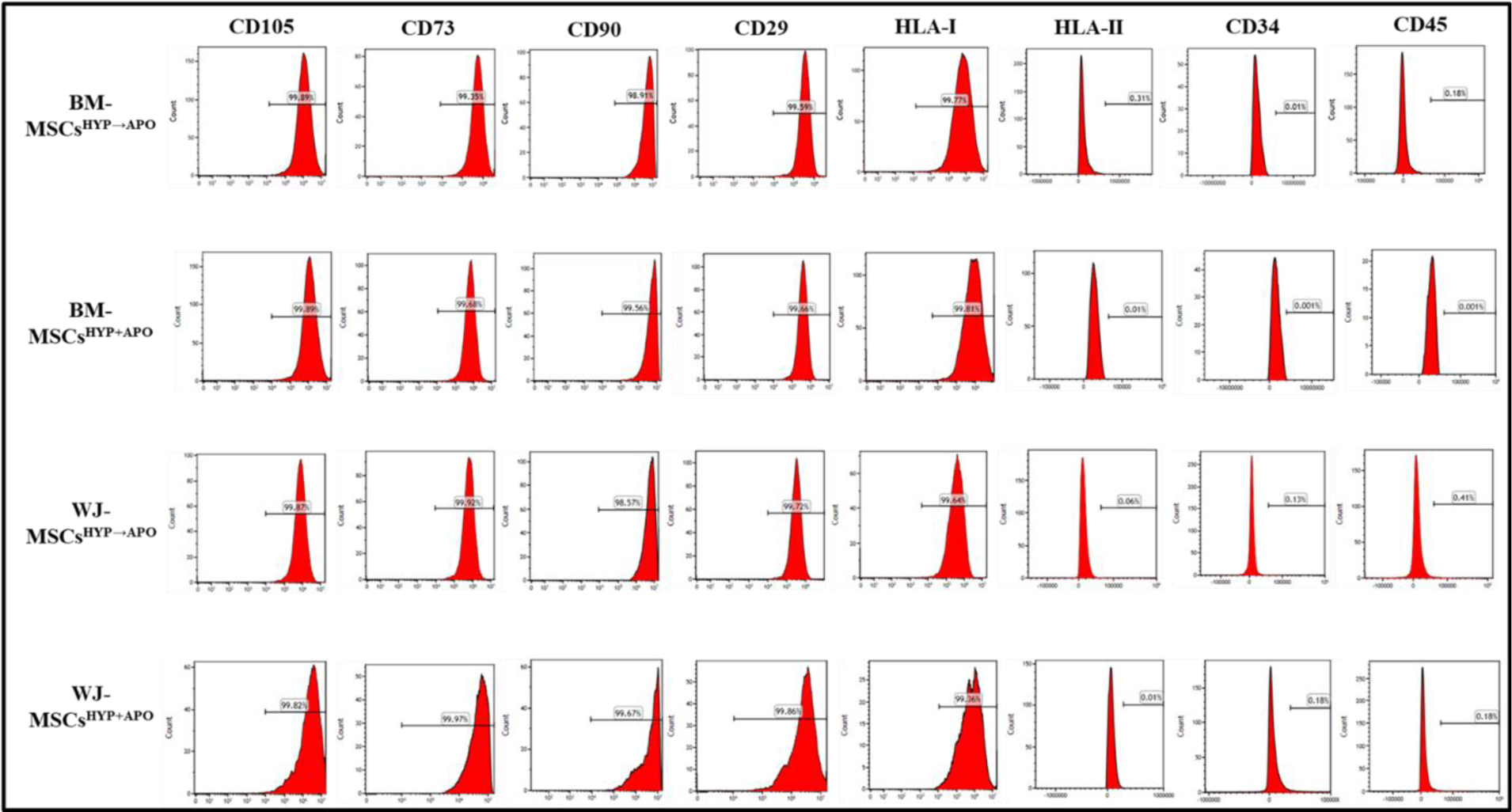
Histograms depicts the surface profile of BM-MSCs^HYP→APO^, BM-MSCs^HYP+APO^, WJ-MSCs^HYP→APO^, WJ-MSCs^HYP+APO^ using flow cytometry. *Abbreviations: BM: Bone marrow; WJ: Wharton’s Jelly; MSCs: Mesenchymal Stem Cells; HYP: Hypoxia-preconditioned; APO: Apoptosis*.

**Figure S4:**
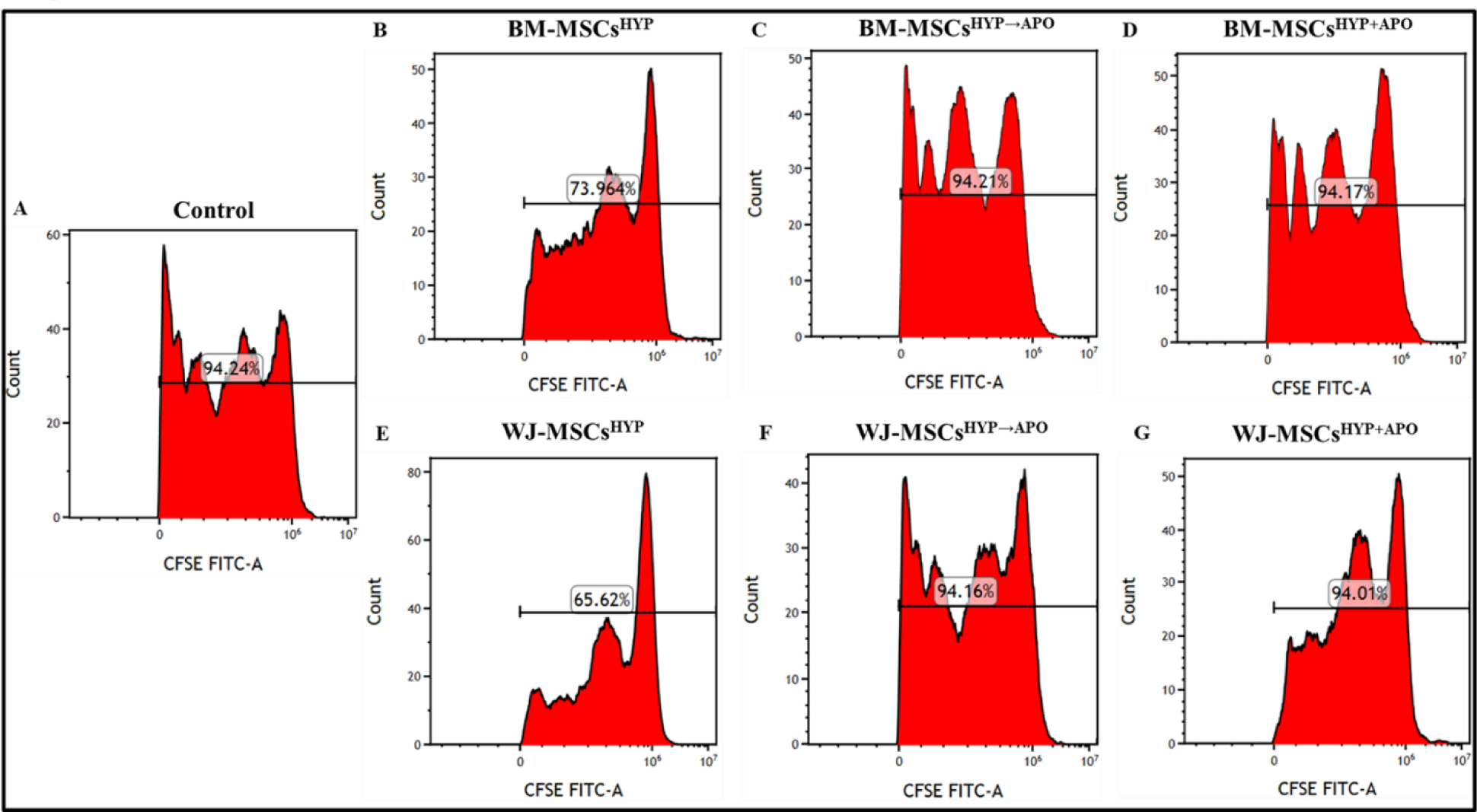
Dot plots showing the proliferation of CD3^+^ T-cell in the direct co-culture of aGVHD patients derived T-cell and **(A)** Untreated (Control). **(B)** BM-MSCs^HYP^. **(C)** BM-MSCs^HYP→APO^. **(D)** BM-MSCs^HYP+APO^. **(E)** WJ-MSCs^HYP^. **(F)** WJ-MSCs^HYP→APO^. **(G)** WJ-MSCs^HYP+APO^. *Abbreviations: BM: Bone marrow; WJ: Wharton’s Jelly; MSCs: Mesenchymal Stem Cells; HYP: Hypoxia-preconditioned; APO: Apoptosis*.

**Figure S5:**
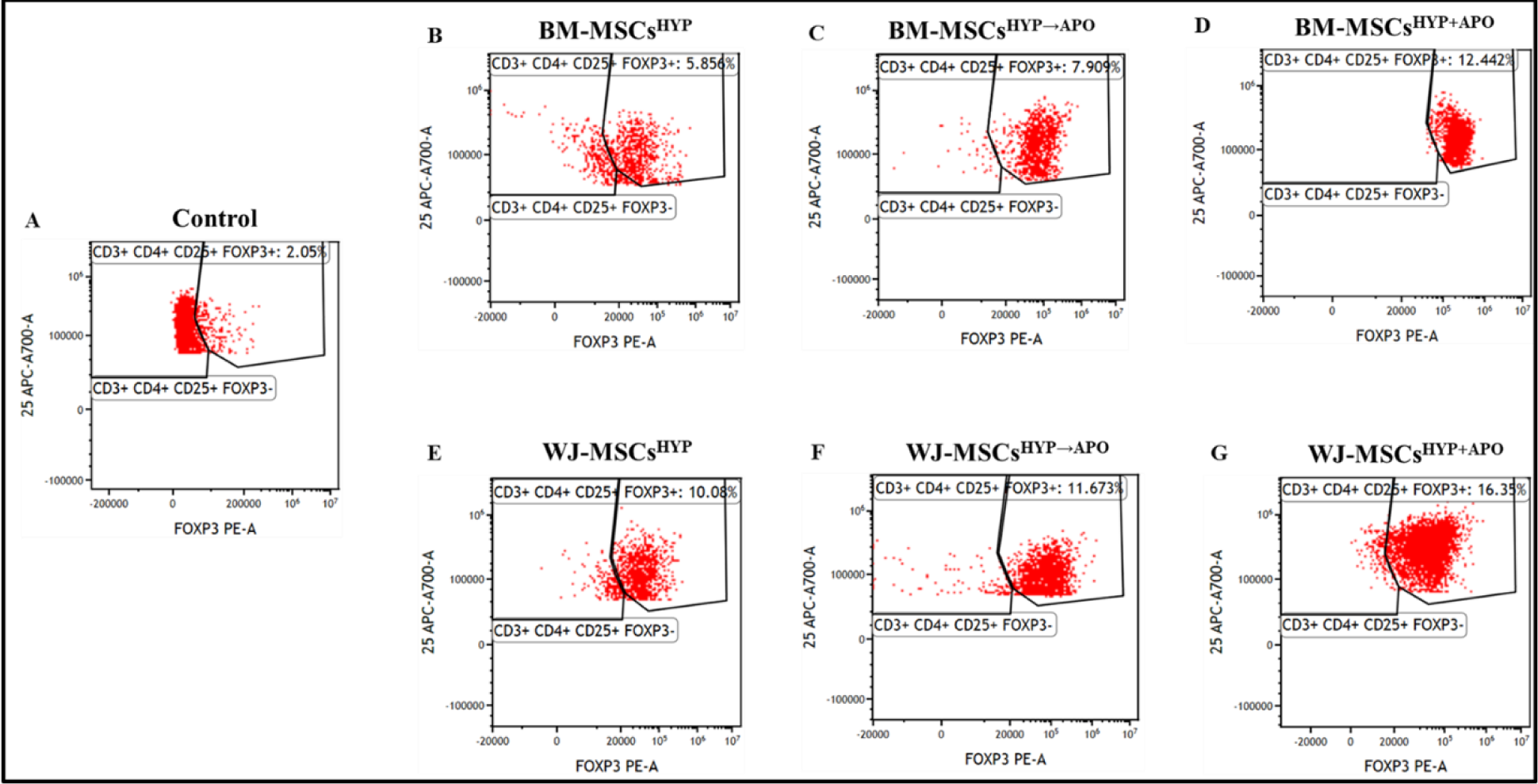
Dot plots showing the percentage of CD3^+^ CD4^+^ CD25^+^ FOXP3^+^ in the direct co-culture of aGVHD patients derived CD3^+^ T-cell and **(A)** Untreated (Control). **(B)** BM-MSCs^HYP^. **(C)** BM-MSCs^HYP→APO^. **(D)** BM-MSCs^HYP+APO^. **(E)** WJ-MSCs^HYP^. **(F)** WJ-MSCs^HYP→APO^. **(G)** WJ-MSCs^HYP+APO^. *Abbreviations: BM: Bone marrow; WJ: Wharton’s Jelly; MSCs: Mesenchymal Stem Cells; HYP: Hypoxia-preconditioned; APO: Apoptosis*.

**Figure S6:**
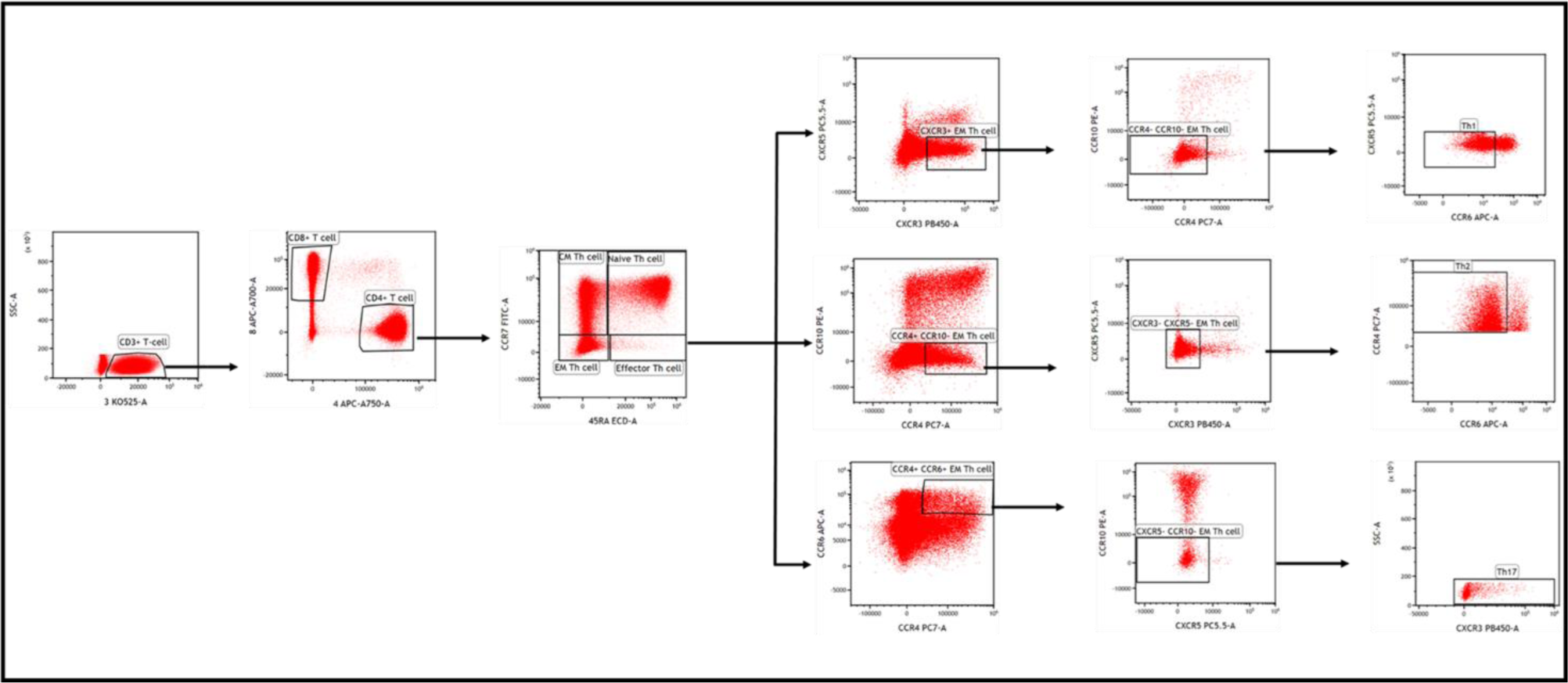
Dot plots showing the gating strategy employed for the enumeration of Th1, Th2, Th17 in the direct co-culture of aGVHD patients derived T-cell and MSCs. *Abbreviations: Th: Helper T cell; EM: Effector Memory; CM: Central Memory*.

**Figure S7:**
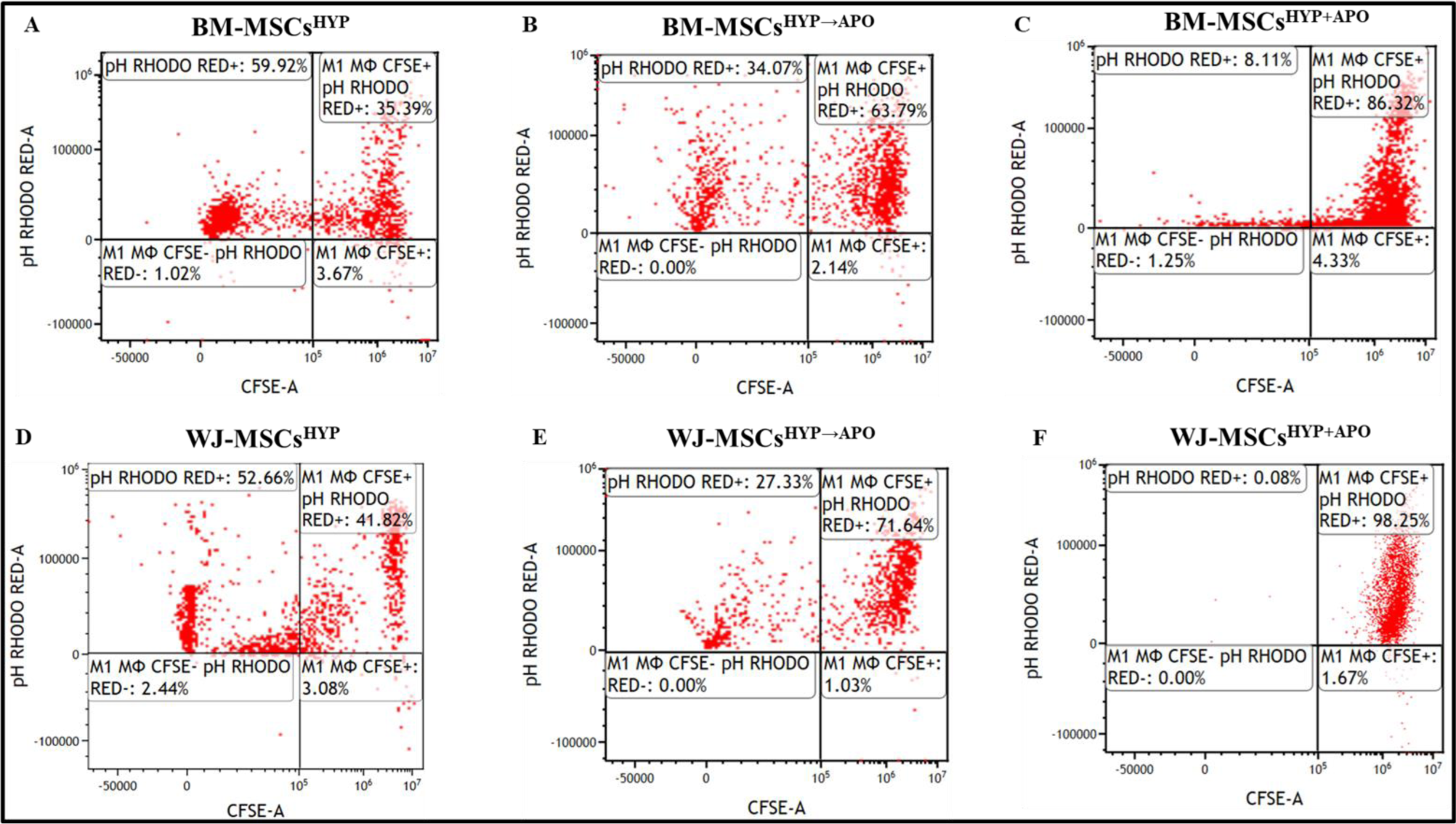
Dot plots showing the efferocytosis of **(A)** BM-MSCs^HYP^ **(B)** BM-MSCs^HYP→APO^ **(C)** BM-MSCs^HYP+APO^ **(D)** WJ-MSCs^HYP^ **(E)** WJ-MSCs^HYP→APO^ **(F)** WJ-MSCs^HYP+APO^ in the co-culture of aGVHD patients derived CD14^+^ monocytes and MSCs. *Abbreviations: BM: Bone marrow; WJ: Wharton’s Jelly; MSCs: Mesenchymal Stem Cells; HYP: Hypoxia-preconditioned; APO: Apoptosis*

**Figure S8:**
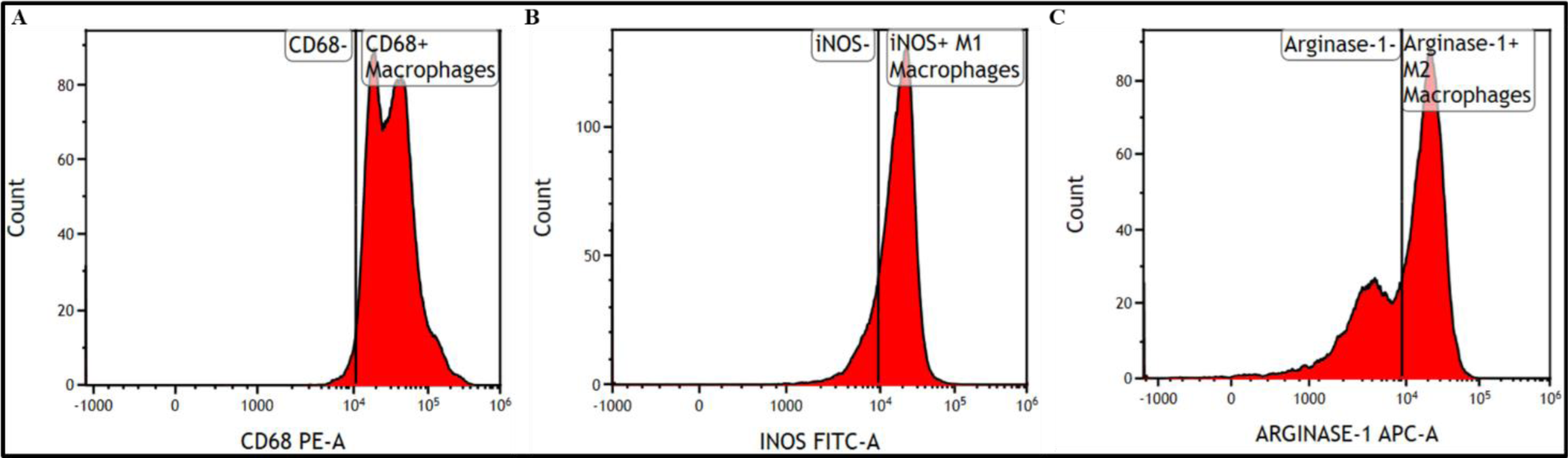
Histogram plots showing the expression of (A) CD68 (B) iNOS (C) Arginase-1 in the co-culture of aGVHD patients derived CD14^+^ monocytes and MSCs.

## Supplementary Table

**Table S1:**
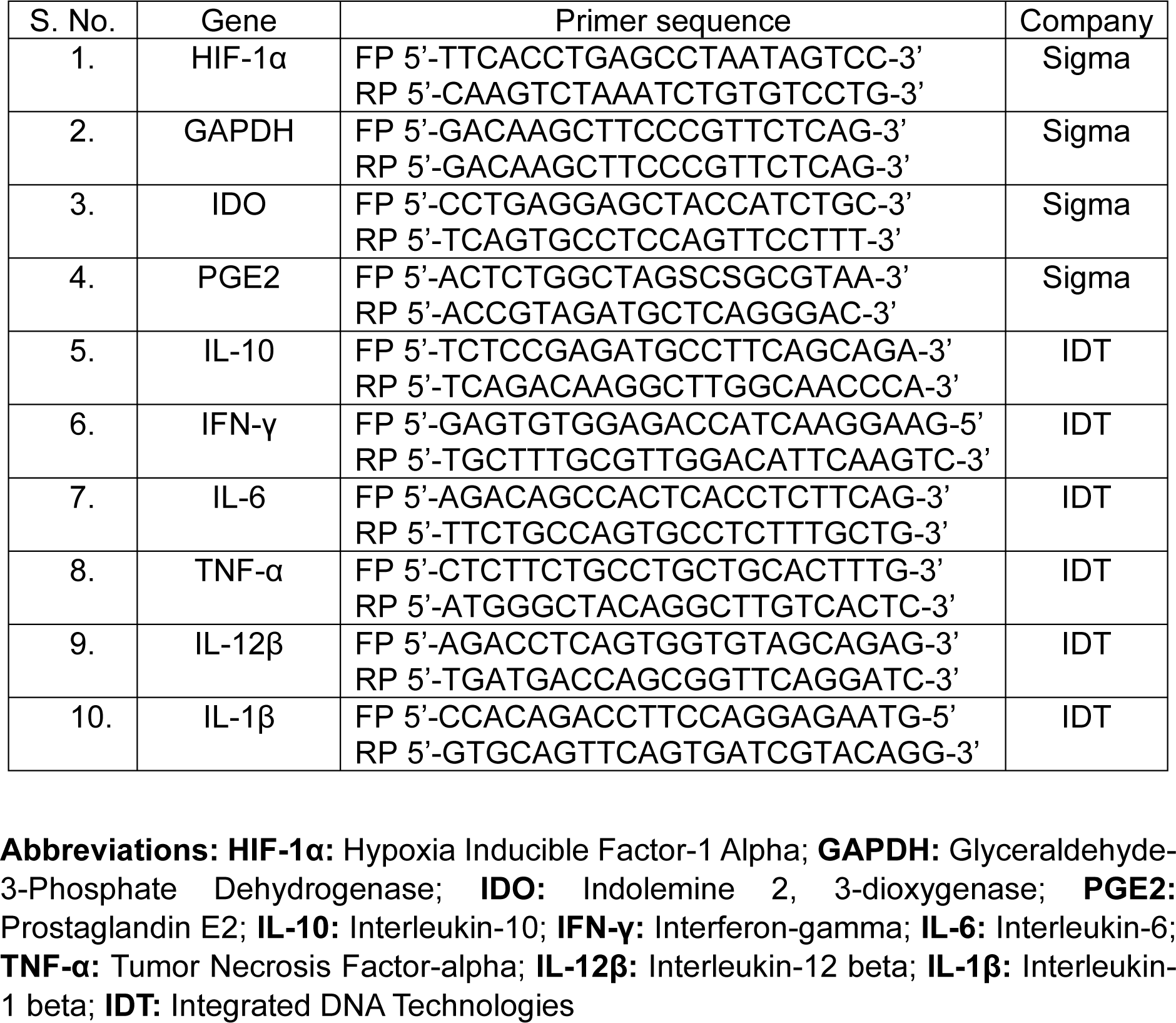
List of primers used to analyze gene expression using qPCR.

